# Shear-Induced Macrophage Secretome Promotes Endothelial Permeability

**DOI:** 10.1101/2025.06.20.660831

**Authors:** Elysa Jui, Griffin Kingsley, Sarah E. Jimenez, Hong T. Kim Phan, George I. Ezeokeke, Fariha N. Ahmad, Ravi K. Birla, Sundeep G. Keswani, K. Jane Grande-Allen

**Affiliations:** Department of Bioengineering, Rice University, Houston, Texas, USA; Division of Pediatric Surgery, Texas Children’s Hospital, Houston, Texas, USA

## Abstract

**Background:** Discrete subaortic stenosis (DSS) is a pediatric cardiovascular disease marked by fibrotic growth within the left ventricular outflow tract (LVOT), leading to severe complications, including left ventricular hypertrophy, aortic regurgitation, and arrhythmias. Despite surgical intervention, a 20-30% recurrence rate suggests a complex underlying pathophysiology. Elevated flow and resultant hemodynamic shear stress within the LVOT are key factors in DSS development. While effects of shear stress on endothelial cells have been studied, the impact on macrophages and their interactions with endothelial cells remains unclear.

**Methods:** In this study, human monocyte-derived macrophages (MDMs) and human aortic endothelial cells (HAECs) were subjected to shear using a cone-and-plate viscometer. Cellular crosstalk was evaluated through conditioned media (CM) transfers. Gene expression, permeability and chemotaxis assays, immunofluorescent staining, and ELISAs assessed cellular responses.

**Results:** MDMs exposed to shear stress exhibited a pro-inflammatory response with upregulated TNF and CXCL8 genes. HAECs exposed to MDM-CM showed increased expression of inflammatory markers (VCAM-1, ICAM-1) and decreased VE-Cadherin and CD31, indicating increased permeability. Permeability assays confirmed that HAECs became more permeable when exposed to MDM-CM. Chemotaxis assays showed time-dependent monocyte migration in both MDM-CM and HAEC-CM. Immunofluorescent staining revealed diminished VE-Cadherin and CD31 in HAECs exposed to MDM-CM.

**Conclusions:** Overall, pathological shear stress induced macrophages to secrete factors that increased endothelial permeability and perpetuated an inflammatory response. This interaction likely exacerbates fibrosis in DSS, promoting recurrence post-surgery. Understanding these mechanisms opens potential therapeutic avenues targeting inflammatory crosstalk between macrophages and endothelial cells, which could mitigate fibrosis and improve patient outcomes.

## Introduction

Discrete subaortic stenosis (DSS) is a critical pediatric cardiovascular disease, distinguished by fibrotic membrane growth within the left ventricular outflow tract (LVOT).^1^ This condition can lead to severe complications, such as left ventricular hypertrophy, aortic regurgitation, arrhythmias, and potentially fatal outcomes if left untreated. The primary intervention, surgical resection, poses challenges with a recurrence rate of 20-30%, suggesting a complex underlying pathophysiology that remains poorly understood.^1^ This recurrence underscores an aggressive phenotype and highlights a pivotal gap in our understanding of its pathogenesis.

Elevated flow and resultant hemodynamic shear stress within the LVOT, exacerbated by structural heart defects, have been identified as key factors in the development of DSS.^1,2^ While the focus has traditionally been on the direct impacts of shear stress on endothelial cells, the role of macrophages in this dynamic environment has been notably overlooked. After the surgical removal of the fibrotic growth, macrophages within the underlying tissue are exposed to pathological shear stress. The consequences of this exposure and its effects on surrounding cells remain largely unexplored, although we have recently shown that macrophages experience a pro-inflammatory response under such conditions.^3^

In this study, we hypothesized that the pathological shear stress characteristic of DSS causes pro-inflammatory macrophages to disrupt the endothelial lining, resulting in increased endothelial permeability. This change facilitates the recruitment and infiltration of monocytes, subsequently promoting fibrosis and contributing to the recurrence of fibrotic membrane growth post-surgery. More broadly, vascular permeability is linked to chronic inflammation and the onset of diseases such as atherosclerosis.^4–6^ In atherosclerosis, activated endothelium from pro-inflammatory cytokines promotes extravasation of inflammatory cells, contributing to subsequent fibrosis and sustained plaque development.^4^ Furthermore, recent evidence is accumulating demonstrating the pathogenic role of vascular permeability in the development of pulmonary fibrosis.^7–9^

We used a cone-and-plate viscometer bioreactor to apply shear stress to human monocyte-derived macrophages (MDMs) and human aortic endothelial cells (HAECs) and evaluated cellular crosstalk through conditioned media (CM) transfers between the two cell types. By investigating the interaction between macrophages and endothelial cells under these conditions, this research aimed to illuminate a previously underexplored aspect of DSS pathophysiology, opening new avenues for therapeutic strategies aiming not only to treat DSS but also to prevent its recurrence, significantly impacting patient outcomes and quality of life.

## Materials and Methods

### Primary human monocyte-derived macrophages isolation and culture

Human monocytes were isolated from donor buffy coats obtained from the Gulf Coast Regional Blood Center (Houston, TX) using density gradient centrifugation. All blood samples were from male donors. Use of de-identified blood product samples received through a third party were deemed by the Rice University IRB to not be human subjects research. Briefly, buffy coats were layered onto Histopaque (1.077 g/mL; Sigma-Aldrich, St. Louis, MO) and centrifuged at 400 x g for 30 minutes at room temperature without brake. Peripheral blood mononuclear cells (PBMCs) were collected from the interphase, pooled with 2-3 other donors, and washed twice with 5 mM EDTA in 1x PBS. PBMCs were then cultured in RPMI 1640 (Sigma-Aldrich) supplemented with 10% fetal bovine serum (FBS) and 1% penicillin/streptomycin). After 24 hours, floating lymphocytes were aspirated, and the flasks were washed twice with 1x PBS, leaving the attached monocytes. The monocytes were cultured in an incubator at 37°C with 5% CO2 for 5-7 additional days in RPMI 1640 containing 10% FBS, 1% penicillin/streptomycin, and macrophage-colony stimulating factor (M-CSF; 20 ng/mL; Novus Biologicals, Centennial, CO) to differentiate into MDMs. Media was replenished every other day. At the end of the 5-7 day incubation period, MDMs were harvested via a 15-minute incubation with Accutase (Sigma-Aldrich) at 37°C followed by gentle cell scraping.

### Human aortic endothelial cell culture

The human aortic endothelial cells (HAECs) used in this study were purchased from Lonza (Walkersville, MD). HAECs were cultured in an incubator at 37°C with 5% CO2 using EGM-2 (Lonza) supplemented with 1% penicillin/streptomycin instead of the gentamicin provided in the bullet kit. The culture media was replenished every other day. HAECs were used at passages less than 6.

### Shear stress application and conditioned media collection

Uniform laminar shear stress was applied separately to HAECs and MDMs using a cone-and-plate viscometer system as previously described.^10–12^ The cone was placed on a 60 mm petri dish seeded with a monolayer of HAECs, which was then placed on a controllable magnetic stir plate inside an incubator at 37°C with 5% CO2. Prior to shear exposure, each dish of cells was washed with 1x PBS and received 3 mL of fresh EGM-2 media. The cones were rotated at 280 RPM or 660 RPM, corresponding to a shear stress of 15 dynes/cm^2^ and 35 dynes/cm^2^, respectively, according to the governing wall shear stress equation:

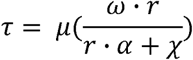

where τ is the shear stress, μ is the dynamic viscosity of the media (0.95 centistokes), ω is the angular velocity of the cone, r is the radius of the cone (30 mm), α is the angle of the cone (0.5°), and χ is the gap height between the cone and the cells (300 µm). HAECs and MDMs were exposed to various magnitudes of shear stress for 3 or 24 hours. Conditioned media (CM) from MDMs and HAECs were then collected and centrifuged at 1000 x g for 5 minutes to remove cellular debris. The supernatants were transferred to new tubes and frozen at −80°C for storage.

### RNA isolation, cDNA synthesis, and quantitative reverse transcription PCR

MDMs and HAECs were seeded into tissue culture-treated 48-well plates and grown to 70% confluency. MDMs and HAECs were serum-starved for 24 hours before exposure to CM. MDMs were exposed to 70% HAEC-CM supplemented with 5% FBS, and HAECs were exposed to 70% MDM-CM supplemented with 5% FBS for 24 hours.

Following CM exposure and direct shear stress exposure, cell phenotypes were analyzed through quantitative reverse transcription PCR (RT-qPCR). RNA was extracted using the Direct-zol™ RNA Microprep kit (Zymo Research, Irvine, CA) according to the manufacturer’s instructions. Briefly, cells were lysed with an equal part by volume TRIzol (ThermoFisher, Waltam, MA) and 95% ethanol mixture. The mixture was then transferred to Zymo-Spin™ IC Columns in a collection tube. Samples were incubated with DNase I at room temperature for 15 minutes. The samples were then washed with RNA wash buffer and then eluted with 15 µL of DNase/RNase-free water. RNA purity (A260/280 > 1.8) was assessed using the NanoDrop 2000 spectrophotometer (ThermoFisher). RNA was stored at −80°C until complementary DNA (cDNA) synthesis.

cDNA was synthesized using the High-Capacity cDNA Reverse Transcription Kit (Applied Biosystems, Waltham, MA) according to manufacturer’s instructions. cDNA samples were stored at −20°C. RT-qPCR was performed on the samples using the iTaq™ Universal Probes Supermix (Bio-Rad Laboratories, Hercules, CA) according to the manufacturer’s instructions. Primers specific to human PECAM1, SNAIL1, VCAM-1, ICAM-1, SELE, ACTA2, COL3A1, VEGF-A, CDH5 and GAPDH were used for HAEC analysis, while primers specific to human CX3CL1, VEGF-A, CCL2, MRC1, TNF, TGFB, CCR7, CXCL8, IL1B, and GAPDH were used for MDM analysis. Relative expression was normalized to GAPDH and the control group. For CM studies, the control group consisted of the cells exposed to CM from the opposite cell type under 3 hours of no shear. For direct shear studies onto HAECs, the control group consisted of the HAECs exposed to 3 hours of no shear.

### Permeability Assay

Endothelial permeability was evaluated using the Endothelial Transwell Permeability Assay Kit (Cell Biologics Inc, Chicago, IL) according to the manufacturer’s instructions. Briefly, HAECs were cultured on a 24-well transwell insert until confluence at 37°C with 5% CO2. Once confluent, HAECs were starved overnight in serum-free media. HAECs were then exposed to MDM-CM for 24 hours. For a negative control, a parallel group of HAECs was exposed to HAEC-CM for 24 hours. MDM-CM and HAEC-CM were then removed and fresh serum-free media with 5 µL streptavidin-HRP was added and incubated for 1 hour at 37°C with 5% CO2. 20 µL of media from the lower chambers were sampled to the wells of a 96-well ELISA assay plate in triplicate. 50 µL of TMB substrate was then added to each well and incubated for 20 minutes at room temperature. Finally, 50 µL of stop solution was added to each well and the absorption values were read at 450 nm on a plate reader. Relative permeability was determined by normalizing to the static 3-hour group.

### Chemotaxis Assay

The ability of MDM-CM and HAEC-CM to induce monocyte migration was assessed using the QCM^TM^ Chemotaxis 5 µm 96-Well Cell Migration Assay (MilliporeSigma, Burlington, MA) according to manufacturer’s instructions. Briefly, MDM-CM and HAEC-CM were added to the wells of the feeder tray. Monocytes were seeded at a density of 20 x 10^4^ cells per well in serum-free media in the migration chamber. The cells were incubated for 24 hours at 37°C with 5% CO2. After incubation, the media was aspirated from the top side of the insert, and the migration chamber plate was placed into a new feeder tray containing prewarmed Cell Detachment Solution and incubated for 30 minutes at 37°C. The medium solution (containing cells that migrated through the membrane into the medium in the feeder tray) and the Cell Detachment Solution (containing cells that migrated through the membrane and were detached by the solution) were combined into a new 96-well plate. Cells were then lysed in a 1:75 solution of CyQuant GR Dye and Lysis Buffer and incubated for 15 minutes. The fluorescence was measured at 480/520 nm using the SpectraMax M2 (Molecular Devices, San Jose, CA). Relative chemotaxis was determined by normalizing to the static 3-hour group.

### Cell viability

Following conditioned media exposure, MDM and HAEC viability was assessed by incubating each sample in a solution containing 2 µM calcein AM and 4 µM ethidium homodimer-1 in PBS (Live/Dead Viability/Cytotoxicity Kit; ThermoFisher) for 30 minutes at room temperature. Samples were imaged immediately via the Eclipse Ti2 fluorescent microscope (Nikon, Melville, NY). Cell viability was quantified through ImageJ/FIJI (NIH).

### Immunofluorescent staining

Following conditioned media exposure, HAECs were stained for CD31, vWF, VE-Cadherin, and DAPI. Briefly, cells were fixed in 4% paraformaldehyde for 20 minutes, permeabilized with 0.5% Triton X for 10 minutes, and blocked in 5% bovine serum albumin (BSA) for 1 hour. Next, cells were incubated with primary antibodies against CD31 (1:200, Invitrogen, Waltham, MA), vWF (1:100, Abcam, Cambridge, UK), or VE-Cadherin (1:100, Invitrogen) overnight at 4°C under gentle agitation. After incubation, cells were washed 3x with 1x PBS and incubated for 1 hour at room temperature with secondary antibodies AF488/AF532 (1:100, Invitrogen). Finally, cells were washed with PBS, counterstained with Hoechst (1:2000, Invitrogen), and imaged using the Eclipse Ti2 fluorescent microscope.

### ELISAs

HAEC-CM from each condition was tested for the presence of endothelial nitric oxide synthase (eNOS) and vWF. eNOS and vWF concentrations were measured using a human eNOS/NOS3 ELISA kit (Invitrogen) or human vWF ELISA kit according to manufacturer’s instructions. Standard calibration curves were used to quantify concentrations.

### Statistical analysis

Statistical analysis was performed using GraphPad Prism (GraphPad Software, La Jolla, CA). A two-way analysis of variance (ANOVA) with Tukey’s post hoc test was used to determine statistical significance (p < 0.05) for each set of conditions in this study. Results are presented as mean ± SD. All experiments were repeated in triplicates at a minimum.

## Results

### MDMs exposed to direct shear vs MDMs exposed to HAECs-CM

MDM gene expression was evaluated after exposure to direct shear stress and HAEC-CM. Interestingly, TNF and CXCL8 experienced a greater response to direct shear stress (**Figure 1**) while CCL2 and VEGF-A were more significantly influenced by exposure to HAEC-CM (**Figure 2**).

**Figure 1.**
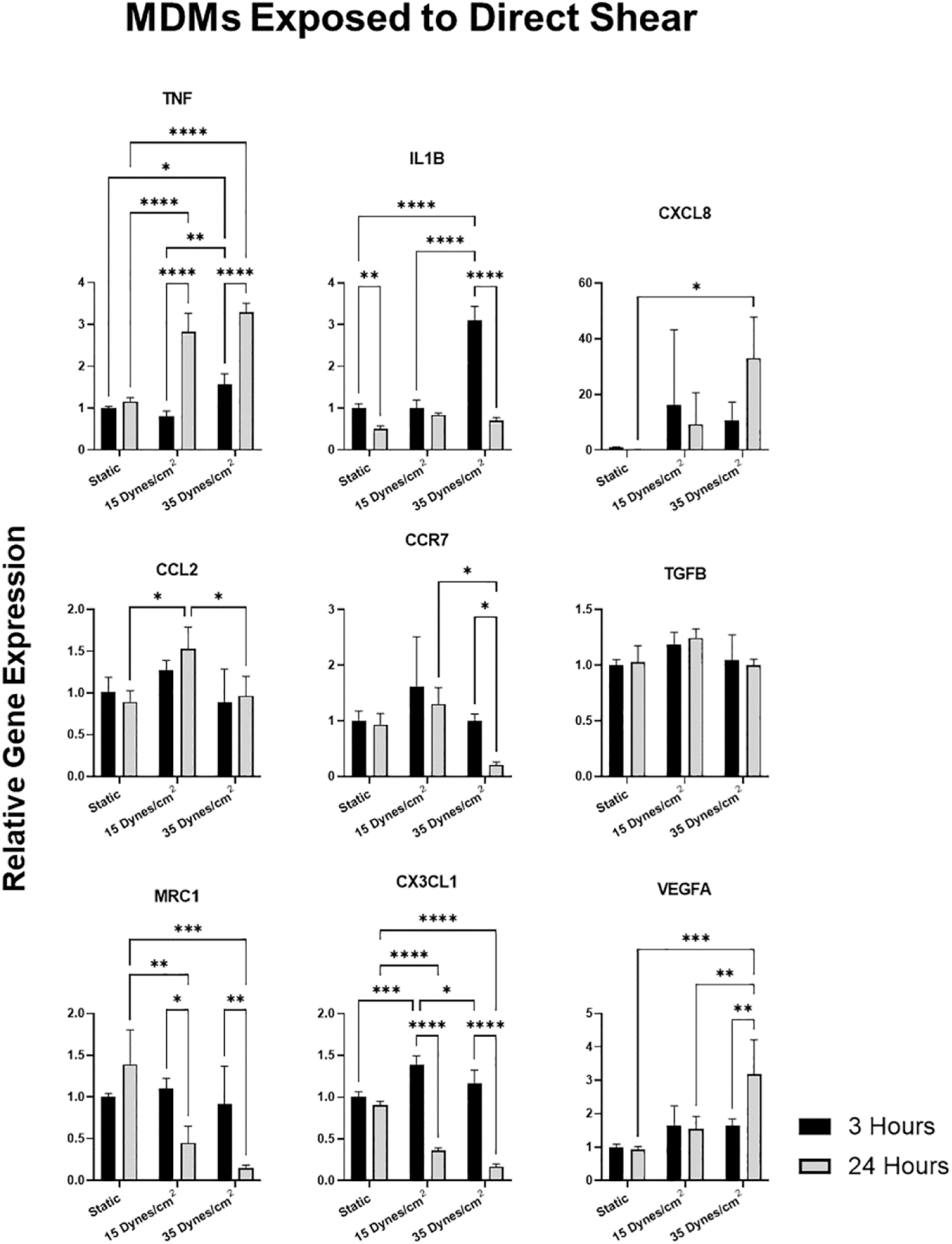
MDM gene expression after exposure to direct shear stress. Gene expression of MDMs exposed to direct shear stress for 3 and 24 hours at 0 dynes/cm^2^ (static), 15 dynes/cm^2^, and 35 dynes/cm^2^. Data is expressed as mean ± SD, n = 3. *p < 0.05, **p < 0.01, ***p < 0.001, ****p < 0.0001. Scale bar = 200 µm.

**Figure 2.**
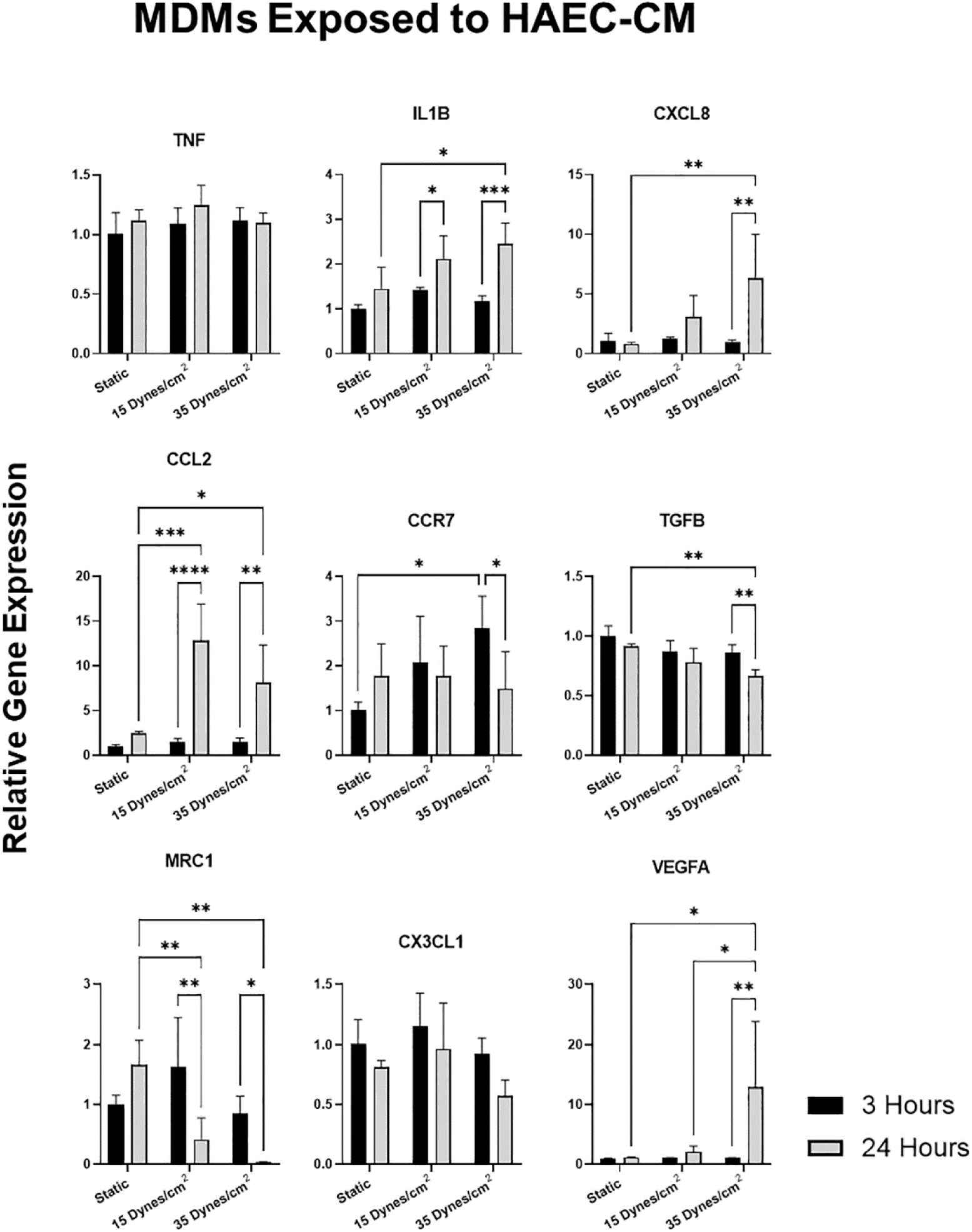
MDM gene expression after exposure to conditioned media. Gene expression of MDMs exposed to conditioned media of HAECs exposed to direct shear stress for 3 and 24 hours at 0 dynes/cm^2^ (static), 15 dynes/cm^2^, and 35 dynes/cm^2^. Data is expressed as mean ± SD, n = 3. *p < 0.05, **p < 0.01, ***p < 0.001, ****p < 0.0001. Legend represents HAECs exposed to shear stress for 3 hours or 24 hours at their respective shear stress conditions.

Specifically, TNF was upregulated at 24 hours in response to direct shear stress, but no significant change in expression levels was observed when exposed to HAEC-CM, indicating a mechanosensitive response. Similarly, CXCL8 exhibited increased expression in response to both direct shear stress (35 dyn/cm^2^) and CM exposure. However, the expression levels were much greater at 24 hours in response to direct shear stress (9.3 & 33.1 fold-change) compared to CM exposure (3.1 & 6.3 fold-change). Due to the high variability in one of the CXCL8 data sets from direct shear (3 hours, 15 dyn/cm^2^) in **Figure 1**, the CXCL8 data was prepared as a box plot in **Supplemental Figure 1A**. This plot showed that the data set has a low median value with one high value outlier. Due to this variance, and the overlap of the data range with the relevant comparison groups, there was no statistically significant comparison involving that shear magnitude.

CCL2 and VEGF-A also demonstrated similar results at 24 hours in response to both conditions, but the magnitude of response was greater with CM exposure (CCL2: 1.5 & 0.97 fold-change vs 12.8 & 8.1 fold-change) (VEGF-A: 1.5 & 3.2 fold-change vs 12.8 & 8.2 fold-change). Because the VEGF-A data set from the CM exposure condition (24 hours, 35 dyn/cm^2^) had high variability, we also examined the VEGF-A data as a box plot (**Supplemental Figure 1B**). This data set had 2 values that were very similar and one value that was much higher, yet the range of the data, on the whole, did not overlap with that of the comparison groups, so a statistically significant difference was detected. Interestingly, IL1-β was upregulated at 3 hours in response to direct shear stress but was upregulated at 24 hours in response to HAEC-CM exposure. This suggests that HAECs exposed to shear stress may sustain the pro-inflammatory response in macrophages. Importantly, the viability of both MDMs and HAECs were not affected by CM exposure (**Supplemental Figures 2-3**).

### HAECs exhibit a pro-inflammatory phenotype when exposed to MDM-CM

HAEC gene expression was evaluated after exposure to direct shear stress and MDM-CM. Interestingly, the inflammatory genes VCAM-1, ICAM, and E-Selectin were largely unaffected or downregulated by direct shear stress, except for an initial increase in ICAM at the 3-hour timepoint (**Figure 3**). However, when HAECs were exposed to MDM-CM, these genes were upregulated at 24 hours, suggesting that shear stress causes MDMs to promote a pro-inflammatory response in HAECs (**Figure 4**).

**Figure 3.**
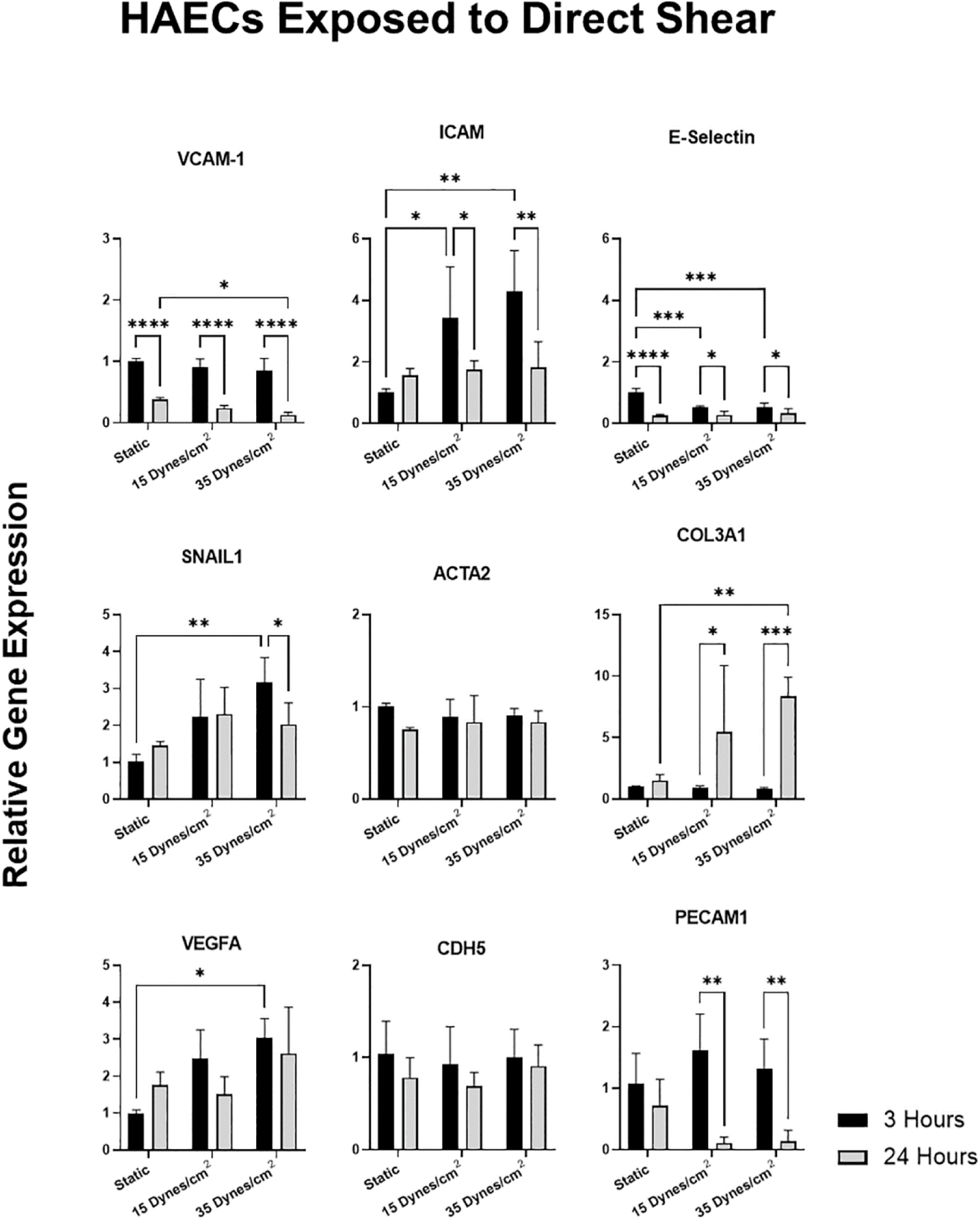
HAEC gene expression after exposure to direct shear stress. Gene expression of HAECs exposed to direct shear stress for 3 and 24 hours at 0 dynes/cm^2^ (static), 15 dynes/cm^2^, and 35 dynes/cm^2^. Data is expressed as mean ± SD, n = 3. *p < 0.05, **p < 0.01, ***p < 0.001, ****p < 0.0001.

**Figure 4.**
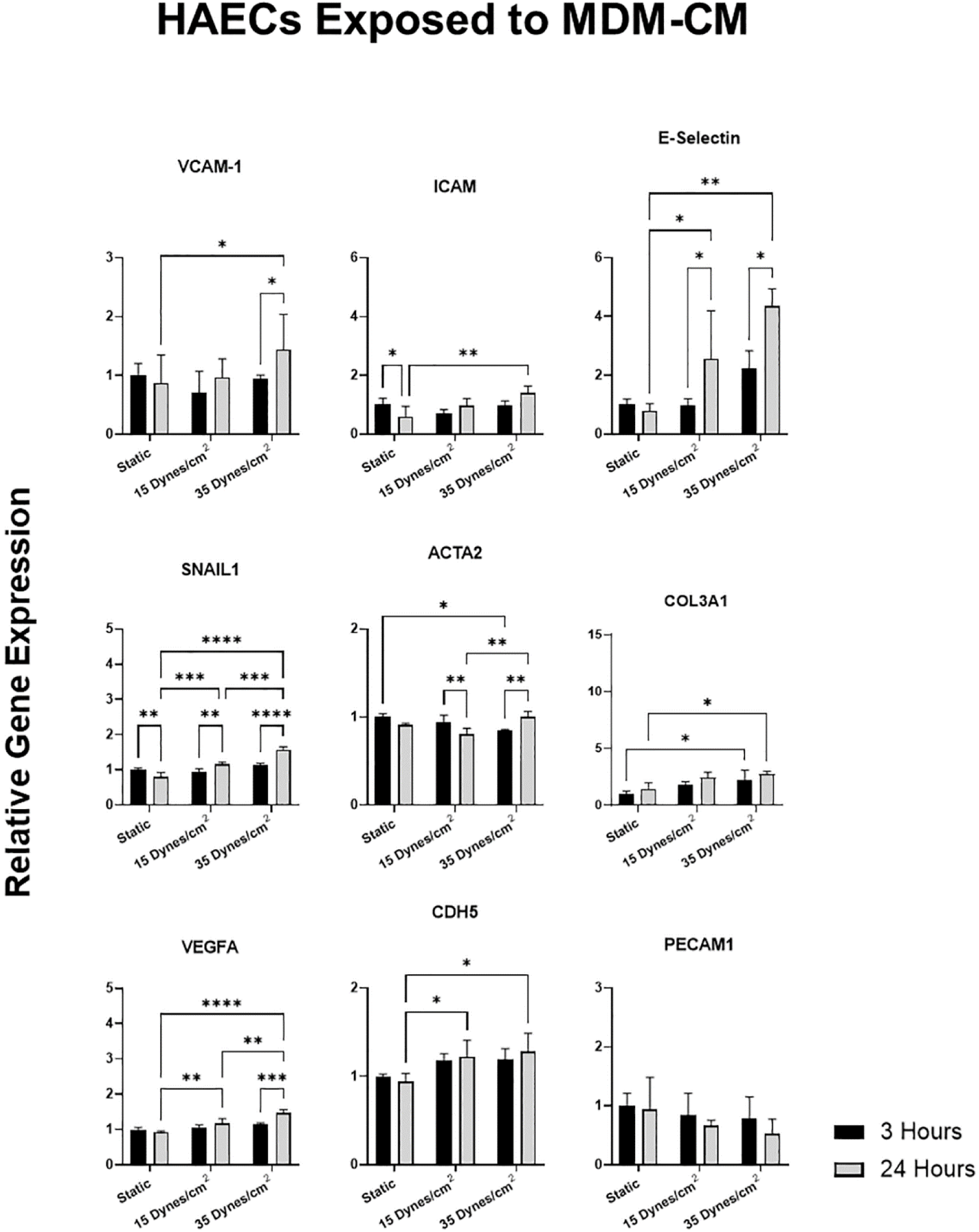
HAEC gene expression after exposure to conditioned media. Gene expression of HAECs exposed to conditioned media of MDMs exposed to direct shear stress for 3 or 24 hours at 0 dynes/cm^2^ (static), 15 dynes/cm^2^, and 35 dynes/cm^2^. Data is expressed as mean ± SD, n = 3. *p < 0.05, **p < 0.01, ***p < 0.001, ****p < 0.0001.

PECAM1 expression remained relatively unchanged in response to both conditions, with a slight but non-significant downregulation observed. EndMT markers SNAIL1 and COL3A1 were upregulated at 3 and 24 hours, respectively, in response to direct shear stress (35 dyn/cm^2^). The COL3A1 expression resulting from 24 hours of direct shear stress (15 dyn/cm^2^) was also high but due to high variability (one high value outlier as shown by the box plot in **Supplemental Figure 1C**), there was no significant difference between that condition and the static loading condition. The expression of E-selectin by the MDM-CM treated HAECs was also highly variable at the 24 hours, 15 dyn/cm^2^ condition. Despite the distribution of values across a large range for this condition, the box plot in **Supplemental Figure 1D** shows that the data ranges for this analysis were non-overlapping so that statistically significant differences could be detected.

When exposed to MDM-CM, SNAIL1 and COL3A1 were also upregulated, but to a lesser extent. Similarly, VEGF-A was upregulated in response to direct shear stress at 3 hours and again at 24 hours in response to MDM-CM, though at a lower magnitude. These results suggest that while direct shear stress is the dominant factor in driving endMT and angiogenesis, MDMs may also contribute to sustaining these processes.

### HAECs experience increased permeability when exposed to MDM-CM

Given the pro-inflammatory phenotype observed in HAECs in response to MDM-CM, we next investigated their permeability, since endothelial cells are known to become more permeable when inflamed.^5,13^ Permeability did not increase when HAECs were exposed to their own CM (negative control) but did increase when exposed to MDM-CM (**Figure 5A**). This result suggests that direct shear stress causes MDMs to secrete cytokines that enhance HAEC permeability, facilitating monocyte infiltration. We then performed a chemotaxis assay comparing the ability of MDM-CM and HAEC-CM to evaluate monocyte infiltration. Interestingly, monocyte chemotaxis was not affected by shear stress and decreased at 24 hours in both MDM-CM and HAEC-CM, indicating a time-dependent response (**Figure 5B**).

**Figure 5.**
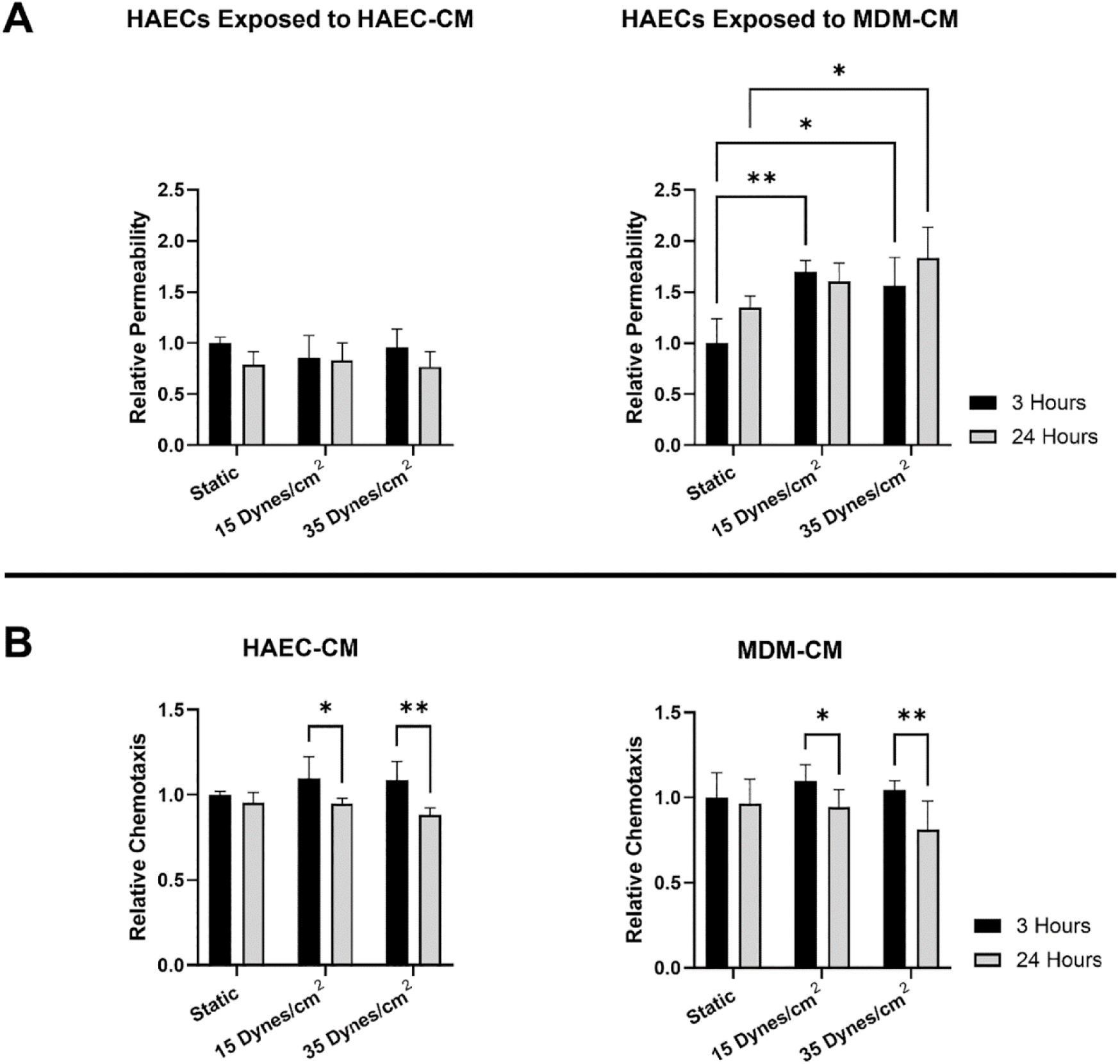
HAEC permeability and CM chemotaxis. (A) Relative permeability of HAECs exposed to their own CM or MDM-CM normalized to the 3-hour static groups. (B) Relative chemotaxis of HAEC-CM and MDM-CM normalized to the 3-hour static groups.

### MDM-CM largely diminishes protein expression of VE-Cadherin, CD31, and vWF

To further evaluate endothelial permeability and function, we exposed HAECs to their own CM or to MDM-CM and stained for VE-Cadherin, CD31, and vWF. VE-Cadherin expression was evidently decreased when HAECs were exposed to MDM-CM compared to their own CM (**Figure 6, Supplemental Figure 4**). This reduction was especially pronounced in the high shear stress condition at 24 hours, where VE-Cadherin was almost completely absent when exposed to MDM-CM, indicating increased permeability with less expression. CD31 expression was also markedly diminished when HAECs were exposed to MDM-CM (**Figure 6, Supplemental Figure 5**). Interestingly, CD31 expression at 3 hours in HAECs exposed to their own CM appeared to decrease in response to shear stress. These outcomes seem to align with the gene expression results. Additionally, vWF expression was also lower in HAECs exposed to MDM-CM (**Figure 6, Supplemental Figure 6**).

**Figure 6.**
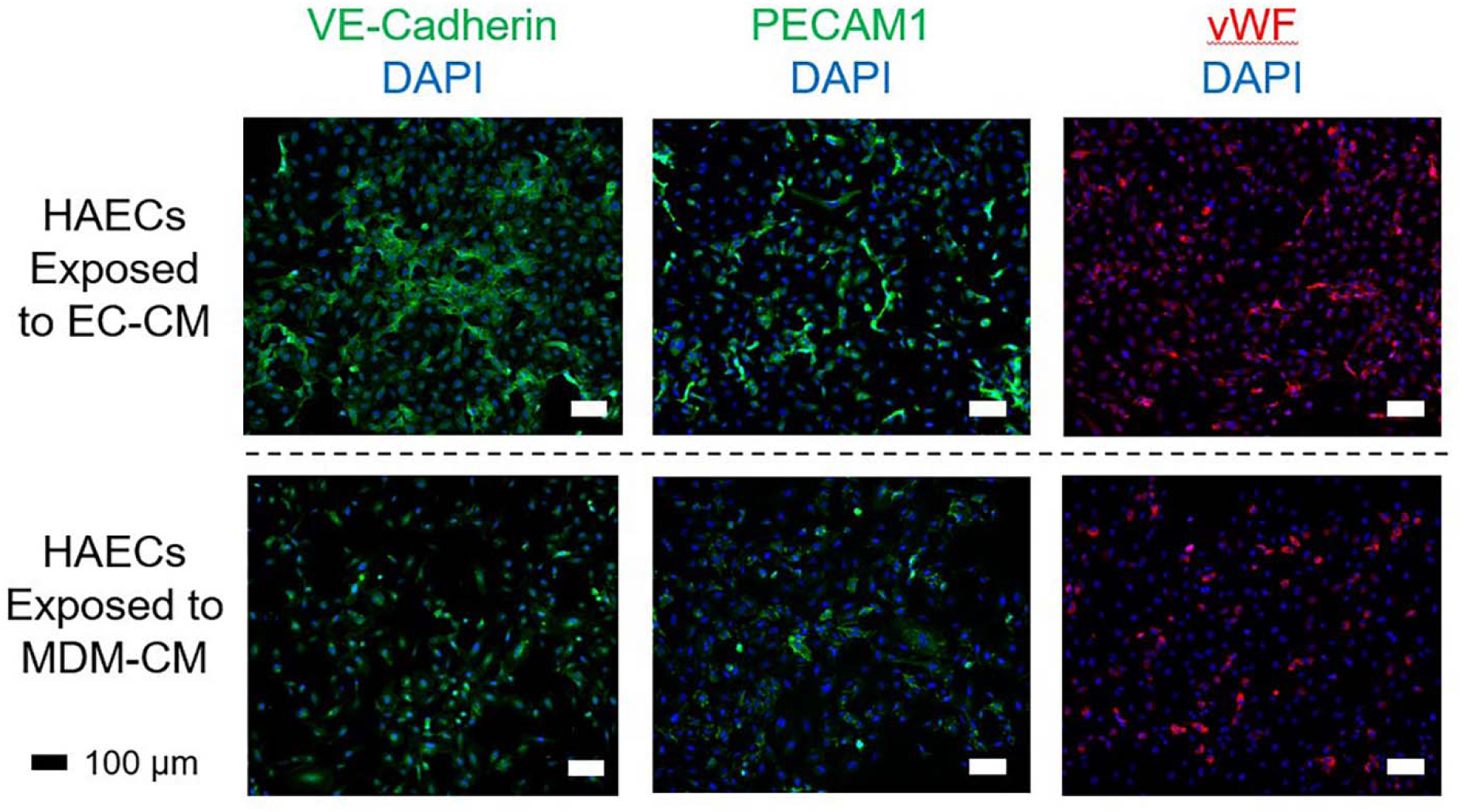
Immunofluorescent staining of HAECs treated with conditioned medium. Immunofluorescent staining of VE-Cadherin (green, column 1), PECAM1 (green, column 2), and vWF (red, column 3) on HAECs after being exposed to their own conditioned medium (row 1) or MDM-conditioned medium (row 2). Conditioned medium was collected from cells exposed to high shear stress (35 dynes/cm^2^) for 24 hours. Scale bar = 100 µm.

### Comment

Overall, we sought to delineate the cellular crosstalk involved in the pathogenesis of the “aggressive” phenotype observed in DSS recurrence. Our results demonstrated that monocyte derived macrophages (MDMs) and HAECs exhibit distinct gene expression profiles in response to direct shear stress and CM. Direct application of shear stress to macrophages resulted in a time-dependent pro-inflammatory response. Specifically, IL-1β gene expression was upregulated at 3 hours, while TNF-α gene expression was upregulated at 24 hours. However, when MDMs were exposed to HAEC-CM, there was an increase in IL-1β gene expression at 24 hours, but no significant change in TNF-α expression. The differential responses observed in TNF and IL-1β gene expression upon exposure to HAEC-CM suggest distinct regulatory mechanisms.

It is important to clarify that our experimental model centers on fully differentiated MDMs rather than undifferentiated monocytes. This distinction is biologically significant because macrophages and monocytes differ in phenotype and function once monocytes exit the circulation and mature within tissues. The mechanosensitivity of monocytes themselves is an important and biologically relevant topic, and previous studies have shown that shear stress can modulate monocyte inflammatory responses;^14^ however, investigating monocyte behavior under shear lies outside the framework of the current work and would require a separate set of experiments. Our model was specifically designed to mimic the mechanical environment within the left ventricular outflow tract (LVOT) following discrete subaortic stenosis (DSS) resection,^3^ where macrophages, whether resident or recruited, would encounter altered flow conditions directly. Within this context, our findings demonstrate that shear-activated macrophages drive endothelial dysfunction and adhesion molecule expression, which we interpret as contributing to fibrotic recurrence in DSS. While the hypothesis that specific monocyte subsets may be preferentially mechanosensitive is intriguing and potentially relevant in other vascular contexts, it was not directly tested or required for the inflammatory amplification mechanism we describe here. Future studies may explore whether specific monocyte populations exhibit shear-responsive phenotypes upstream of the macrophage behaviors characterized in this work.

NF-κB activation, typically responsible for TNF regulation,^15^ might be less influenced by endothelial-derived factors compared to direct shear stress. This suggests that HAECs, when exposed to shear stress, secrete factors that selectively influence specific inflammatory pathways in MDMs. The increase in IL-1β expression indicates that HAEC-derived factors may activate the NLRP3 inflammasome pathway in MDMs, which can be activated independently of NF-κB. This selective activation highlights the complex interplay between HAECs and MDMs in the inflammatory response. Investigating these pathways in greater detail could uncover specific therapeutic targets to modulate inflammatory responses more precisely.

Direct shear stress on HAECs promoted endMT, a phenomenon that is well-established in the literature.^16,17^ However, the exposure of HAECs to MDM-CM not only sustained this response, as seen through the upregulation of endMT markers SNAIL1 and COL3A1, but also induced a pronounced pro-inflammatory phenotype, characterized by increased expression of adhesion molecules (VCAM-1, ICAM, E-Selectin). This suggests that macrophages, when activated by shear stress, can induce endothelial dysfunction and promote processes such as endMT, which are critical in fibrosis and tissue remodeling.

The response of VCAM-1, ICAM-1, and E-Selectin in HAECs to direct shear stress warrants further discussion. While laminar shear ultimately enforces an anti-inflammatory endothelial state, the temporal dynamics of these responses are important to consider. At the onset of shear stress, there is a well-documented transient activation of inflammatory signaling, often through NF-κB induction.^18,19^ Sudden exposure of static endothelial cells to high shear can acutely activate integrin-dependent mechanotransduction pathways, in which activated integrins and focal adhesion kinase (FAK) trigger NF-κB signaling within minutes.^20^ This rapid NF-κB activation likely explains the initial increase in ICAM-1 we observed at the 3-hour timepoint, as ICAM-1 is a known NF-κB target gene that can be induced by flow onset through these mechanosensitive pathways.^21^ However, unlike sustained disturbed flow, laminar shear does not maintain this proinflammatory signaling. Rather, endothelial cells under prolonged laminar shear adapt and enter a quiescent state, with NF-κB activity dampened by reinstated IκB and rising levels of KLF2/4, which actively refocus gene expression toward a protective program.^22^ Importantly, KLF2 selectively inhibits VCAM-1 and E-Selectin expression, blocking their transcriptional induction even in the presence of inflammatory stimuli, but exerts less restraint on ICAM-1.^23^ This selective inhibition likely explains our observation that VCAM-1 and E-Selectin were largely unaffected or downregulated by shear; any slight NF-κB activation at flow onset is insufficient to overcome the robust suppressive influence of KLF2/KLF4 on these promoters. ICAM-1, by contrast, showed a transient uptick at 3 hours before returning to baseline, consistent with the idea that ICAM-1 is less directly restrained by KLF factors and thus mirrors the brief early NF-κB activation. By 24 hours of continuous flow, the shear-conditioned endothelium expresses lower levels of all three adhesion molecules compared to static conditions, with ICAM-1 subsiding and VCAM-1/E-Selectin remaining low. This pattern of early ICAM-1 induction followed by overall downregulation of adhesion molecules under steady shear is consistent with prior endothelial cell studies and provides a mechanistic framework for the responses observed in our direct shear experiments.

Increased permeability of HAECs, confirmed through permeability assays and immunofluorescent staining for VE-Cadherin and CD31, is a critical finding. VE-Cadherin and CD31 are integral to cell-cell adhesion and maintaining endothelial barrier integrity.^13^ Their decreased expression suggests a disruption of endothelial junctions, leading to increased permeability. This further supports the notion that MDMs contribute to endothelial barrier dysfunction, facilitating monocyte infiltration and subsequent fibrotic growth. Although our chemotaxis assays indicated that monocyte migration was not directly affected by shear stress, the increased permeability of endothelial cells would still allow more monocytes to infiltrate the tissue, contributing to the chronic inflammatory milieu in DSS.

Interestingly, CCL2 gene expression in MDMs was modestly upregulated in response to direct shear stress but profoundly increased when exposed to HAEC-CM. CCL2, coding for MCP-1, is a potent monocyte chemoattractant. This could explain why there were no significant changes in chemotaxis observed in MDM-CM due to shear stress since CCL2 was only profoundly upregulated when MDMs were exposed to HAEC-CM. This suggests that shear stress on endothelial cells induces them to secrete cytokines that, in turn, activate macrophages to attract more monocytes. Additionally, CXCL8, which codes for IL-8, a potent neutrophil chemoattractant, was upregulated to a much greater extent than CCL2, indicating that while monocyte chemotaxis is not predominantly driven by shear stress, neutrophil chemotaxis should be investigated further as a potential contributor to the inflammatory response in DSS.

Our findings suggest that pathological shear stress induces macrophages to secrete factors that not only perpetuate an inflammatory response but also increase endothelial permeability. This interaction likely exacerbates the fibrotic response in DSS, promoting the recurrence of fibrotic growth post-surgery. Understanding these mechanisms opens potential therapeutic avenues targeting the inflammatory crosstalk between macrophages and endothelial cells. For instance, anti-inflammatory therapies or cytokine inhibitors could be explored to mitigate fibrosis and improve patient outcomes in DSS.

Future studies should focus on identifying the specific signaling pathways involved in the crosstalk between macrophages and endothelial cells under shear stress. It will be especially relevant to examine this signaling and crosstalk-driven cell behaviors in the setting of inflammatory mediators. Proteomic analyses of HAEC-CM and the use of pathway-specific inhibitors would be useful. Additionally, exploring the temporal dynamics of these interactions *in vivo* will provide deeper insights into DSS progression and aid in the development of more effective treatment strategies. Longitudinal studies in animal models of DSS could validate these findings and assess the therapeutic potential of targeting these inflammatory pathways.

## Acknowledgements

Financial support for this research was provided by a gift from Lew and Laura Moorman, NIH R01 HL140305 (to J-GA and SK), an NSF Graduate Research Fellowship Program (to EJ), an AHA Research Supplement (25DIVSUP1478725 to GE), and funding for undergraduate research interns (SJ, GK) through the American Heart Association (20UFEL35260054) and the NIH MARC program (5T34GM136452). The graphical abstract was prepared using BioRender.

## References

1. Massé DD, Shar JA, Brown KN, Keswani SG, Grande-Allen KJ, Sucosky P. Discrete Subaortic Stenosis: Perspective Roadmap to a Complex Disease. Front Cardiovasc Med. 2018;5:122.

2. Shar JA, Brown KN, Keswani SG, Grande-Allen J, Sucosky P. Impact of Aortoseptal Angle Abnormalities and Discrete Subaortic Stenosis on Left-Ventricular Outflow Tract Hemodynamics: Preliminary Computational Assessment. Front Bioeng Biotechnol. 2020;8:114.

3. Jui E, Kingsley G, Phan HKT, Singampalli KL, Birla RK, Connell JP, Keswani SG, Grande-Allen KJ. Shear Stress Induces a Time-Dependent Inflammatory Response in Human Monocyte-Derived Macrophages. Ann Biomed Eng. 2024;52(11):2932–47.

4. Botts SR, Fish JE, Howe KL. Dysfunctional Vascular Endothelium as a Driver of Atherosclerosis: Emerging Insights Into Pathogenesis and Treatment. Front Pharmacol. 2021;12:787541.

5. Claesson-Welsh L, Dejana E, McDonald DM. Permeability of the Endothelial Barrier: Identifying and Reconciling Controversies. Trends Mol Med. 2021;27(4):314–31.

6. Mundi S, Massaro M, Scoditti E, Carluccio MA, van Hinsbergh VWM, Iruela-Arispe ML, De Caterina R. Endothelial permeability, LDL deposition, and cardiovascular risk factors-a review. Cardiovasc Res. 2018;114(1):35–52.

7. Jayant G, Kuperberg S, Somnay K, Wadgaonkar R. The Role of Sphingolipids in Regulating Vascular Permeability in Idiopathic Pulmonary Fibrosis. Biomedicines. 2023;11(6).

8. May J, Mitchell JA, Jenkins RG. Beyond epithelial damage: vascular and endothelial contributions to idiopathic pulmonary fibrosis. J Clin Invest. 2023;133(18).

9. Probst CK, Montesi SB, Medoff BD, Shea BS, Knipe RS. Vascular permeability in the fibrotic lung. Eur Respir J. 2020;56(1).

10. Heo KS, Lee H, Nigro P, Thomas T, Le NT, Chang E, McClain C, Reinhart-King CA, King MR, Berk BC, Fujiwara K, Woo CH, Abe J. PKCζ mediates disturbed flow-induced endothelial apoptosis via p53 SUMOylation. J Cell Biol. 2011;193(5):867–84.

11. Jo H, Song H, Mowbray A. Role of NADPH oxidases in disturbed flow- and BMP4- induced inflammation and atherosclerosis. Antioxid Redox Signal. 2006;8(9-10):1609–19.

12. Reinhart-King CA, Fujiwara K, Berk BC. Physiologic stress-mediated signaling in the endothelium. Methods Enzymol. 2008;443:25–44.

13. Sukriti S, Tauseef M, Yazbeck P, Mehta D. Mechanisms regulating endothelial permeability. Pulm Circ. 2014;4(4):535–51.

14. Son H, Choi HS, Baek SE, Kim YH, Hur J, Han JH, Moon JH, Lee GS, Park SG, Woo CH, Eo SK, Yoon S, Kim BS, Lee D, Kim K. Shear stress induces monocyte/macrophage-mediated inflammation by upregulating cell-surface expression of heat shock proteins. Biomed Pharmacother. 2023;161:114566.

15. Parameswaran N, Patial S. Tumor necrosis factor-α signaling in macrophages. Crit Rev Eukaryot Gene Expr. 2010;20(2):87–103.

16. Islam S, Boström KI, Di Carlo D, Simmons CA, Tintut Y, Yao Y, Hsu JJ. The Mechanobiology of Endothelial-to-Mesenchymal Transition in Cardiovascular Disease. Front Physiol. 2021;12:734215.

17. Mahmoud MM, Serbanovic-Canic J, Feng S, Souilhol C, Xing R, Hsiao S, Mammoto A, Chen J, Ariaans M, Francis SE, Van der Heiden K, Ridger V, Evans PC. Shear stress induces endothelial-to-mesenchymal transition via the transcription factor Snail. Sci Rep. 2017;7(1):3375.

18. Mohan S, Mohan N, Sprague EA. Differential activation of NF-kappa B in human aortic endothelial cells conditioned to specific flow environments. Am J Physiol. 1997;273(2 Pt 1):C572–8.

19. Orr AW, Stockton R, Simmers MB, Sanders JM, Sarembock IJ, Blackman BR, Schwartz MA. Matrix-specific p21-activated kinase activation regulates vascular permeability in atherogenesis. J Cell Biol. 2007;176(5):719–27.

20. Petzold T, Orr AW, Hahn C, Jhaveri KA, Parsons JT, Schwartz MA. Focal adhesion kinase modulates activation of NF-kappaB by flow in endothelial cells. Am J Physiol Cell Physiol. 2009;297(4):C814–22.

21. Imberti B, Morigi M, Zoja C, Angioletti S, Abbate M, Remuzzi A, Remuzzi G. Shear stress-induced cytoskeleton rearrangement mediates NF-kappaB-dependent endothelial expression of ICAM-1. Microvasc Res. 2000;60(2):182–8.

22. Marin T, Gongol B, Chen Z, Woo B, Subramaniam S, Chien S, Shyy JY. Mechanosensitive microRNAs-role in endothelial responses to shear stress and redox state. Free Radic Biol Med. 2013;64:61–8.

23. SenBanerjee S, Lin Z, Atkins GB, Greif DM, Rao RM, Kumar A, Feinberg MW, Chen Z, Simon DI, Luscinskas FW, Michel TM, Gimbrone MA, Jr., García-Cardeña G, Jain MK. KLF2 Is a novel transcriptional regulator of endothelial proinflammatory activation. J Exp Med. 2004;199(10):1305–15.

