## Supplemental Material for "Shear-Induced Macrophage Secretome Promotes Endothelial Permeability"

**SUPPLEMENTAL FIGURES**

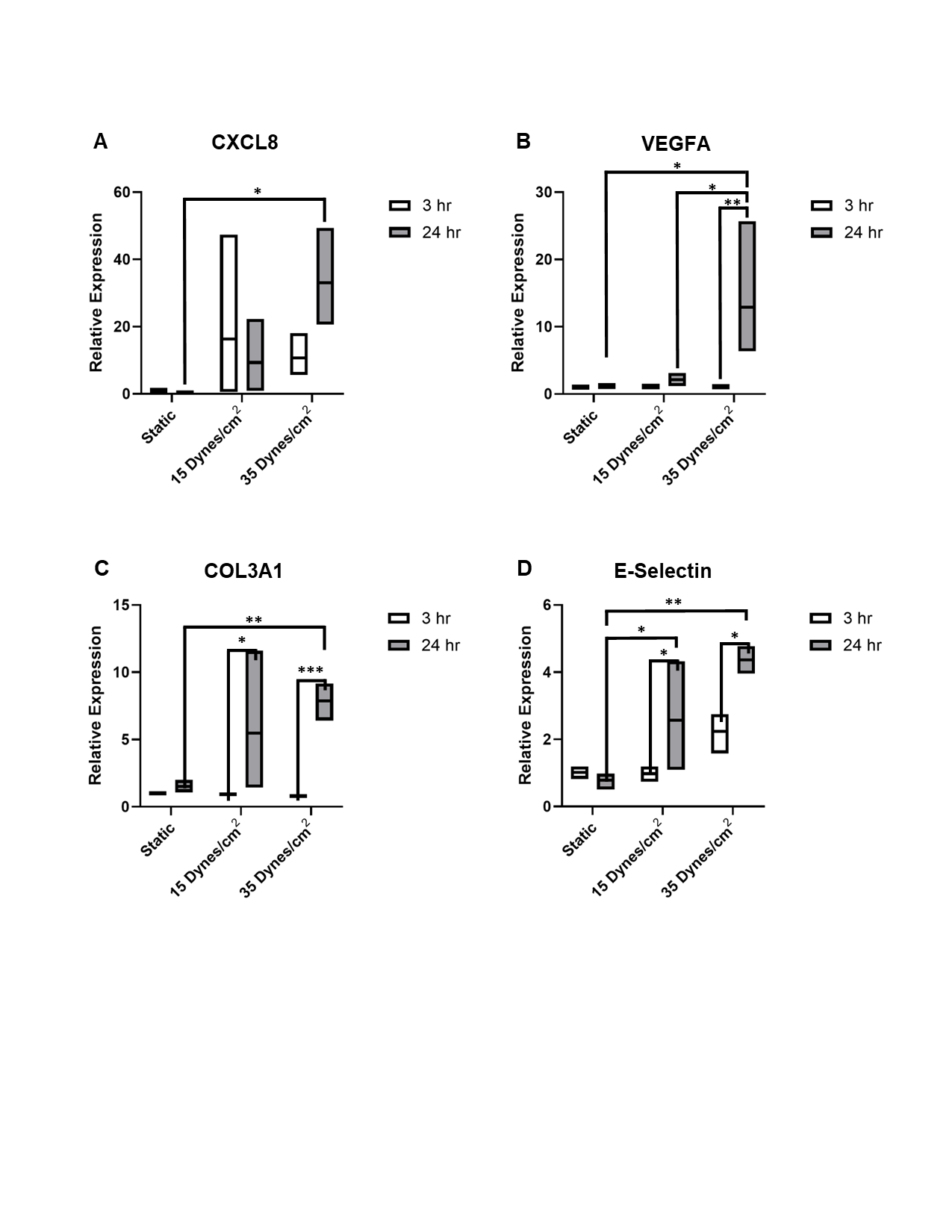

**Supplemental Figure 1. Selected data sets presented as box plots to evaluate variability.**

Gene expression of cells exposed to either direct shear or CM for 3 hours or 24 hours. Box plots show the median, 25^th^, and 75^th^ percentiles of the data range (n = 3). (A) CXCL8 expression from MDMs exposed to direct shear. (B) VEGF-A expression from MDMs exposed to HAEC-CM. (C) COL3A1 expression from HAECs exposed to direct shear. (D) E-Selectin expression from HAECs exposed to MDM-CM.

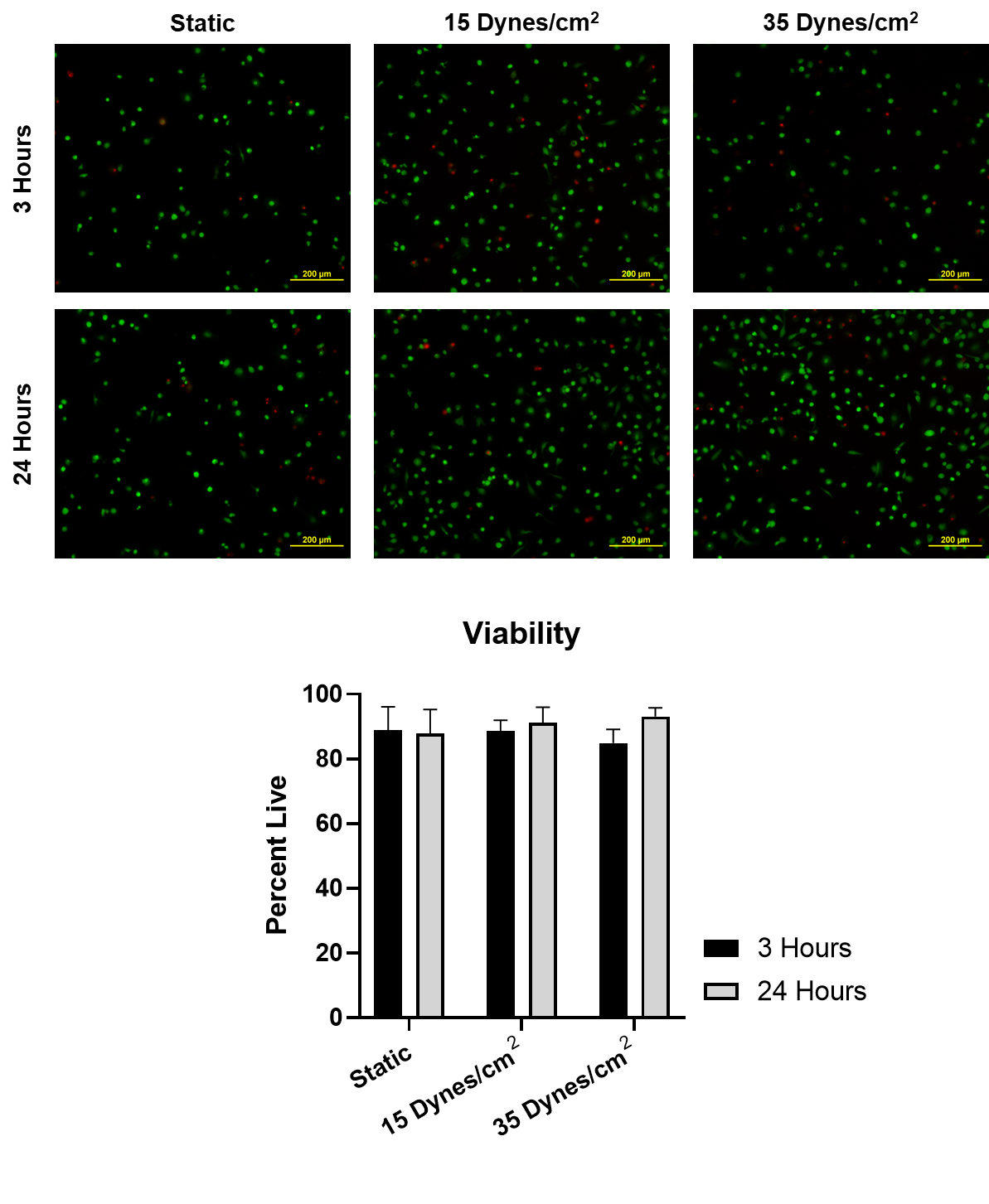

**Supplemental Figure 2. MDM viability after exposure to conditioned media.**

Live/dead staining and quantification of MDMs exposed to HAEC-CM. Data is expressed as mean ± SD, n = 3. Legend represents HAECs exposed to shear stress for 3 hours or 24 hours at their respective shear stress conditions.

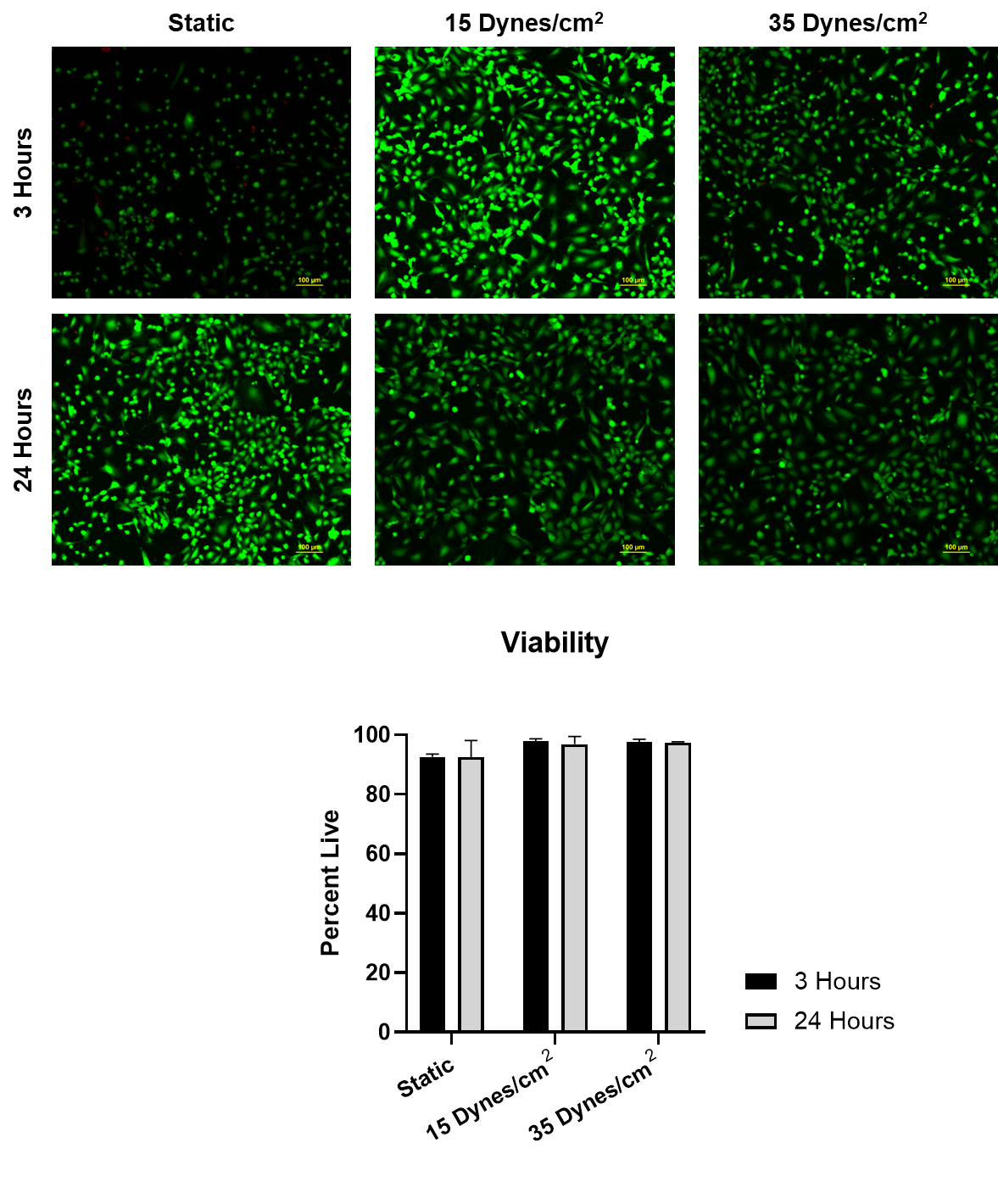

**Supplemental Figure 3. HAEC viability after exposure to conditioned media.**

Live/dead staining and quantification of HAECs exposed to MDM-CM. Legend represents MDMs exposed to shear stress for 3 hours or 24 hours at their respective shear stress conditions. Data is expressed as mean ± SD, n = 3. Scale bar = 100 µm.

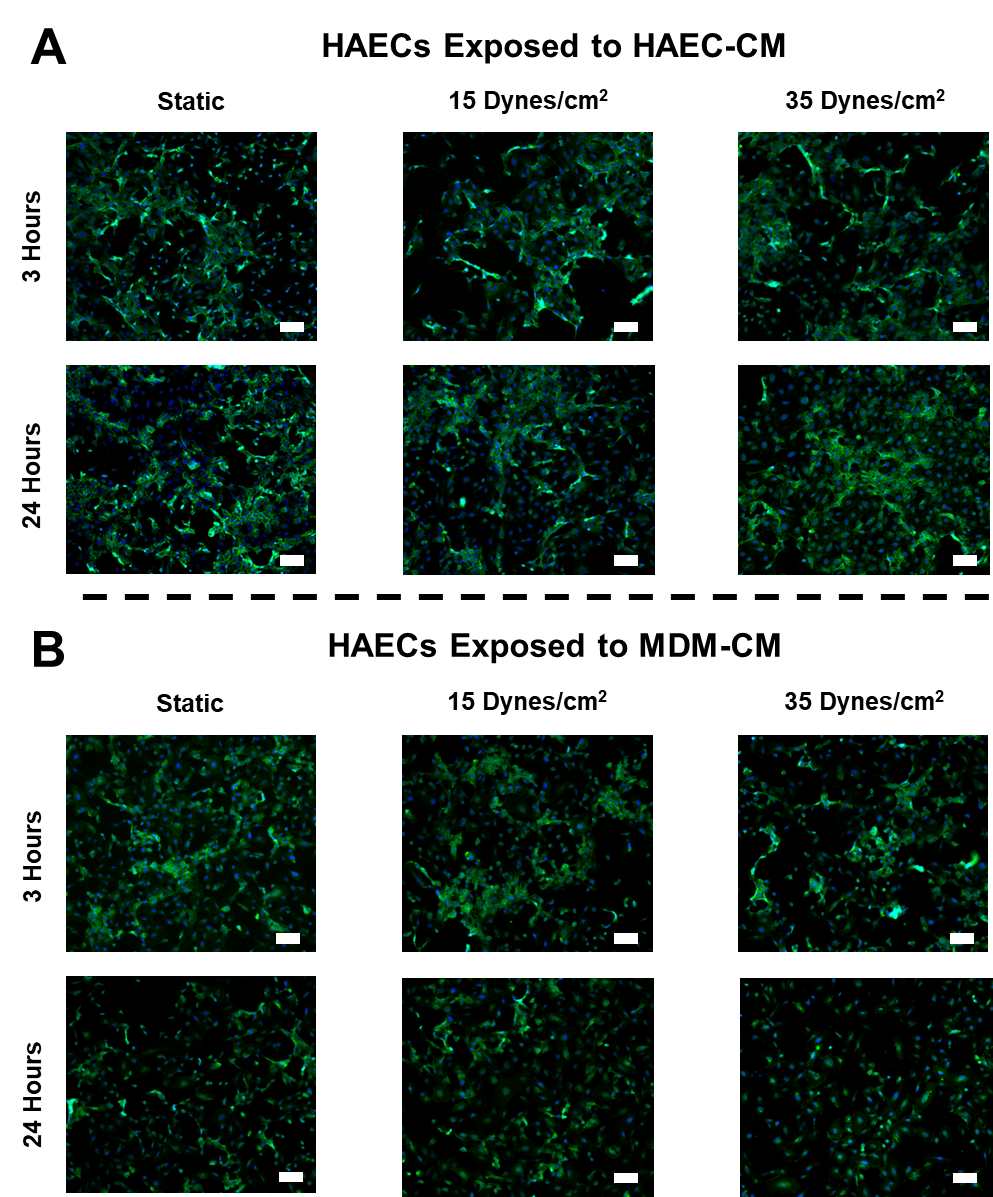

**Supplemental Figure 4. Immunofluorescent staining of VE-Cadherin.**

Immunofluorescent staining of VE-Cadherin (green) on HAECs after being exposed to (A) their own CM or (B) MDM-CM. Scale bar = 100 µm.

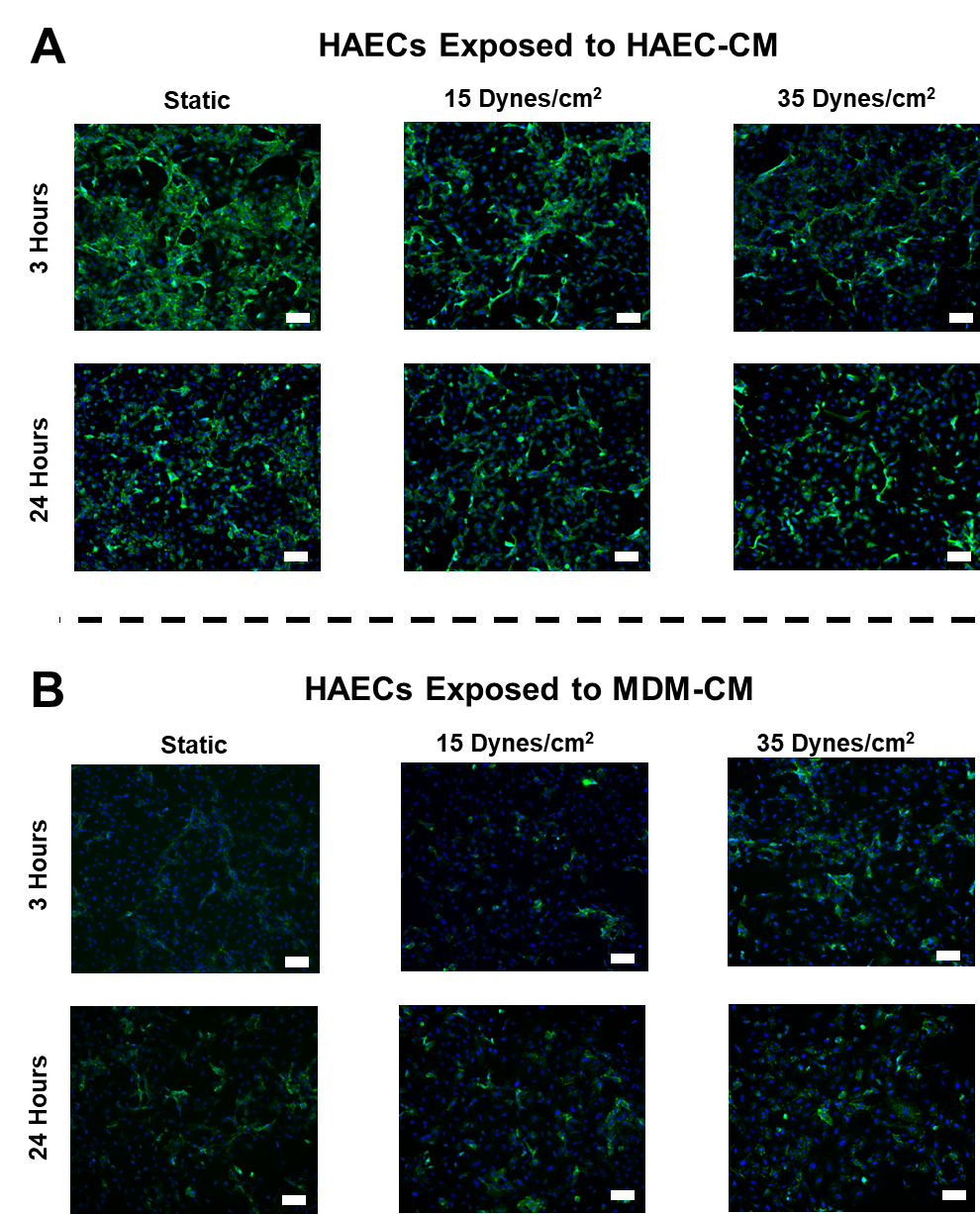

**Supplemental Figure 5. Immunofluorescent staining of PECAM-1.**

Immunofluorescent staining of PECAM-1 on HAECs after being exposed to (A) their own CM or (B) MDM-CM. Scale bar = 100 µm.

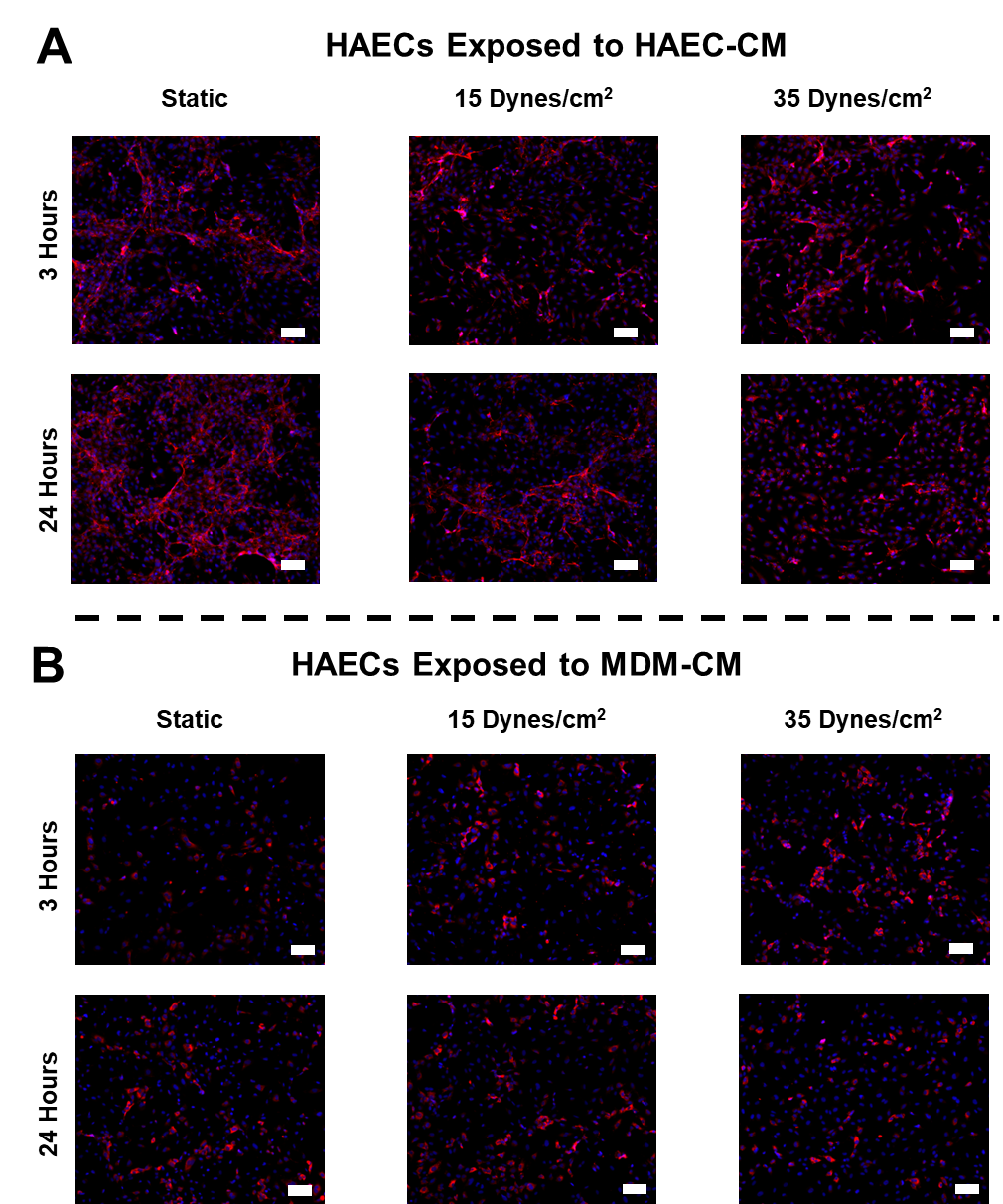

**Supplemental Figure 6. Immunofluorescent staining of vWF.**

Immunofluorescent staining of vWF on HAECs after being exposed to (A) their own CM or (B) MDM-CM. Scale bar = 100 µm.
